# The Isl1/Shh/Wnt5a cascade controls the invasion of myoblasts by tuning the CXCL12/CXCR4 axis during tongue morphogenesis

**DOI:** 10.1101/2021.10.25.465629

**Authors:** Wei Zhang, Jiaojiao Yu, Guoquan Fu, Ruiqi Huang, Jianying Li, Huarong Huang, Jing Liu, Dongliang Yu, Lin Gan, Mengsheng Qiu, Zunyi Zhang, Feixue Li

## Abstract

The migration and invasion of myoblast cells derive from the occipital somites is critical for tongue morphogenesis. However, the molecular mechanisms underlying myoblast migration and invasion remain elusive. In this study, we report that transcription factor Isl1 is required for tongue development. Loss of Isl1 in the distal mandibular epithelium results in aglossia due to the defect of myoblast cells invasion. In the absence of Isl1 expression, myoblast cells in the first branchial arch fail to migrate into the presumptive tongue primordium. Down-regulation of Shh was detected in the distal mandibular epithelium after Isl1 deletion. As a downstream target gene of Shh, Wnt5a attracts the migration of mesenchymal cells expressing chemokine ligand 12 (CXCL12). CXCL12^+^ mesenchymal cells in the Isl1 mutant embryo fail to migrate to the distal region but accumulate in a relatively small proximal domain of the first branchial arch. CXCL12 serves as a bidirectional cue for myoblasts that express its receptor CXCR4, attracting CXCR4^+^ myoblasts invasion at low concentration but repelling at high concentration. The defect in the distal migration of CXCL12^+^ mesenchymal cells leads to a high local concentration of CXCL12, thereby preventing the invasion of CXCR4^+^ myoblasts. In addition, transgenic activation of Ihh alleviates the defect in tongue development and rescues the invasion of myoblasts, corroborating the functional involvement of Hedgehog signaling in tongue development. In summary, this study provides the first line of genetic evidence for the Isl1/Shh/Wnt5a cascade to regulate myoblasts invasion by tuning the CXCL12/CXCR4 axis.

## Introduction

Human congenital tongue malformations that occur during embryogenesis, such as aglossia, microglossia, and macroglossia, greatly affect the life quality of individuals (1). The tongue development begins with median lingual swelling of the first branchial arch (2). During this stage, the majority of the tongue anlage is composed of neural crest cell (NCC)-derived mesenchymal cells. Then, lateral lingual swellings form on each side of the median tongue bud. Myoblasts derived from different somites migrate along the hypoglossal cord and reach the first arch at about E10.5 (3-5). Most of the tongue muscles originate from the occipital myoblasts, while the connective tissue and vasculature are derived from NCCs which originate from the dorsal neural tube (6). The myoblast cells are precisely regulated to migrate and invade into the tongue primordium. Although it has been found that epithelial-derived signals control tongue morphogenesis (7, 8). However, the mechanism by which epithelial-derived signals regulate myoblast migration and invasion is still unclear.

Sonic hedgehog (Shh), Wnt5a and CXCL12/CXCR4 axis have been reported to be important for cell migration and invasion during organogenesis (8-10). Removing Hedgehog signaling in NCCs (*Wnt1-Cre;Smo*^*n/c*^) leads to a vestigial tongue, and the myogenic progenitors are absent in the tongue anlage (8). Shh signaling is required cell-autonomously to regulate directional muscle cell migration in the limb (11). Wnt5a pathway underlies outgrowth of multiple structures in the vertebrate embryo, and Wnt5a deficiency leads to abnormal development of tongue (10, 12). It has also been reported that Wnt5a regulates directional cell migration and polarity (13-15). Chemokine CXCL12 provides directional cues for the migration of multiple different cell types, such as germ cells, lymphocytes and muscle progenitor cells (16-20). Previous reports indicate CXCL12/CXCR4 regulated the migration of both proliferating and terminally differentiated muscle cells (21). However, it remains unclear how these signaling pathways interact each other between the epithelium, mesenchyme and myoblasts to coordinate the development of the tongue.

Islet1 (Isl1), a member of LIM homeodomain family of transcription factors, has been proved to play critical roles during pattern formation and cell type specification in many developmental process (22, 23). We previously found that Isl1 is exclusively expressed in the distal epithelium of first branchial arch as early as E9.5, and Isl1 ablation (*Isl1*^*shhCre*^) led to the mandibular outgrowth defects (24). In the present study, we reported that the *Isl1*^*shhCre*^ mice exhibited aglossia as a result of defective distal invasion of myoblasts. We further revealed that Isl1/Shh/Wnt5a axis regulates the distal migration of CXCL12^+^ mesenchymal cells, which is essential for establishing the concentration gradient of CXCL12 to attract myoblast invasion. CXCL12 regulates the invasion of CXCR4^+^ myoblasts in a concentration-dependent bidirectional manner. Since the migration and invasion of myoblasts play a consequential role in the development of many organs, the novel mechanism we discovered is of great significance for understanding the mechanism of organogenesis and malformations.

## Results

### Functional Isl1 is required for initiation of tongue

To investigate the potential function of *Isl1* in the development of tongue, we generated mice with epithelial cell specific deletion of *Isl1* by crossing *Shh-Cre* (Fig. S1) with *Isl1*^*fl*^ mice. Scanning electron microscopy (SEM) analysis demonstrated the tongue was absent in *Isl1*^*shhCre*^ embryos (100% penetration) (Fig. 1A, Fig. S2). The expression of *Isl1* in the mandible was determined by immunofluorescence (Fig. 1B) and the *Isl1-LacZ* knock-in allele (Fig. 1C). We demonstrated that the expression of Isl1 was detected in distal ectoderm of the first branchial arch as early as E9.5 (Fig. 1B). Immunohistochemical analysis showed no ISL1 protein expression in *Isl1*^*shhCre*^ mice at E9.5, compared to ISL1 expression in the *Shh*^*Cre*^ control mice (Fig. 1B). Anti-MHC staining revealed that only a stack of residual muscle cells existed in the tongue forming region of the mutant embryos (Fig. 1D). Taste bud was not detected in the *Isl1*^*shhCre*^ embryos using an antibody against Cytokeratin-8 (CK8, Troma-1) (Fig. 1E). Together, these results suggested that an intrinsic function of *Isl1* is required for the tongue formation.

**Figure 1.**
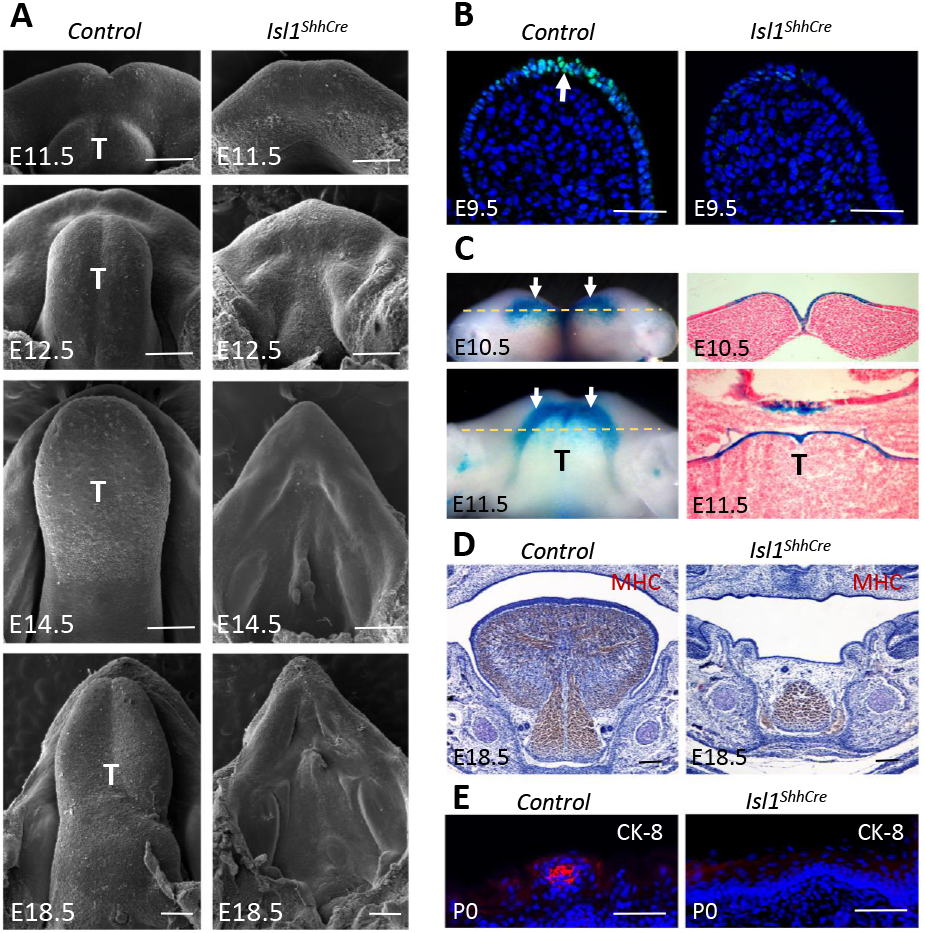
Isl1 is required for mouse tongue morphogenesis. (A) SEM showing the early development of the mouse tongue in *Shh*^*Cre*^ (Control) and *Isl1*^*ShhCre*^ mice. (B) Immunofluorescence staining with Isl1 antibody on the sagittal sections of E9.5 mandibles in control (arrows) and *Isl1*^*ShhCre*^ mice. (C) X-Gal staining showing Isl1 expression (arrows) in *Isl1*^*LacZ/+*^ knock-in mice at E10.5 and E11.5 (left panel for oral transverse view and right panel for coronal sections). (D) Immunohistochemitry with MHC antibody on the coronal sections of E18.5 mandibles in *Shh*^*Cre*^ (Control) and *Isl1*^*ShhCre*^ mice. (E) Immunofluorescence staining with Cytokeratin-8 (CK-8) antibody on the coronal sections of P0 tongues in *Shh*^*Cre*^ (Control) and *Isl1*^*ShhCre*^ mice. T, tongue. Scale bars, 200 μm (A, D), 100 μm (B), 20 μm (E).

### Ablation of Isl1 leads to defective myoblast migration and proliferation

The absence of the majority of tongue muscles in the mutant embryos suggests that the aglossia may be caused by defective myoblast migration into tongue primordium. It has been well documented that Pax3 and Desmin are expressed in early myogenic precursor cells migrating from the somites into the branchial arch at around E10 (25, 26). Immunofluorescent staining revealed that both Pax3- and Desmin-labeled myoblast cells reached the floor of the first branchial arch, but did not migrate into the tongue anlage in the mutant embryos (Fig. 2A; Fig. S3), indicating that myoblast cells failed to migrate and invade into the tongue mesenchyme after they arrived at the first branchial arch in Isl1 mutant mice.

**Figure 2.**
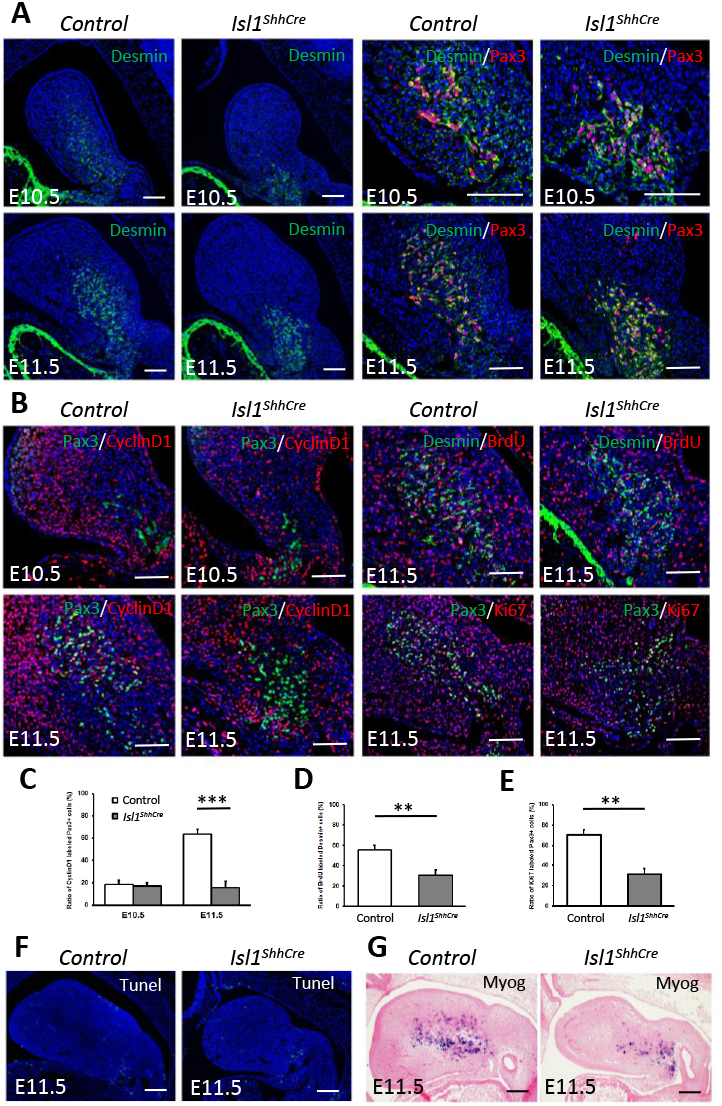
Deletion of Isl1 in the epithelial cells leads to the defect of myoblast cells invasion and proliferation. (A) Double immunofluorescence staining with Desmin and Pax3 antibody on the sagittal sections of E10.5 and E11.5 mandibles. (B) Double immunofluorescence staining with Pax3 and Cyclin D1, Desmin and BrdU, Pax3 and Ki67 on the sagittal sections of E10.5 and E11.5 mandibles. (C) Quantification of Cyclin D1^+^ positive Pax3^+^ cells versus total Pax3^+^ cells in *Shh*^*Cre*^ (Control) and *Isl1*^*ShhCre*^ mice. (D) Quantification of BrdU^+^ positive Desmin^+^ cells versus total Desmin^+^ cells in *Shh*^*Cre*^ (Control) and *Isl1*^*ShhCre*^ mice. (E) Quantification of Ki67^+^ positive Pax3^+^ cells versus total Pax3^+^ cells in *Shh*^*Cre*^ (Control) and *Isl1*^*ShhCre*^ mice. (F) TUNEL staining on the sagittal sections of E11.5 mandibles in *Shh*^*Cre*^ (Control) and *Isl1*^*ShhCre*^ mice. (G) In situ hybridization of Myog on the sagittal sections of E11.5 mandibles in *Shh*^*Cre*^ (Control) and *Isl1*^*ShhCre*^ mice. Error bars represent standard deviations (n=3 to 6 samples). *, P < 0.05. **, P < 0.01. ***, P < 0.001. T, tongue. Scale bars, 200 μm.

We next investigated whether there is a difference in myoblast cell proliferation between the mutant and wild type. Cyclin D1 immunofluorescence showed that the proliferation index of Pax3^+^ myoblasts did not change significantly in the E10.5 mutant, but was markedly slower in the mutant at E11.5 (Fig. 2B, C). Consistently, labeling of BrdU and Ki67 in Desmin^+^ and Pax3^+^ myoblasts were also reduced in the E11.5 mutant (Fig. 2B, D, E). TUNEL analysis revealed no significant change in apoptosis in E11.5 mutant tissues (Fig. 2F). The differentiation of myoblasts was examined by their expression of Myogenin (Myog), a marker for differentiating myoblasts, and no apparent difference was detected in both wild and mutant embryos (Fig. 2G). These results demonstrated that in the mutant embryo, the proliferation of myoblasts was reduced after reaching the first branchial arch, but their differentiation and cell survival was not affected.

### Loss of Isl1 results in decreased expression of Shh in distal ectoderm of the first branchial arch

To determine how epithelial localized Isl1 affects the migration of muscle cells, RNA-seq was used to identify the downstream target genes of Isl1 in the first branchial arch epithelium. Of 15,340 genes analyzed, 399 protein-coding genes were differentially expressed by ≥ 1.5-fold in the *Isl1*^*Shh-Cre*^ mutant versus the control (Table S1). Among them, 178 genes were down-regulated and 221 genes were up-regulated (Table S1). Gene ontology (GO) term analyses revealed that the significantly affected biological processes including multicellular organism development, cell fate commitment, regulation of transcription, somitogenesis, etc. were enriched (Fig. 3B). The expression of *Shh*, which plays a key role in these biological processes, was significantly down-regulated in the mutant (Fig. 3C, Table S1), suggesting that the Hedgehog signal activity in the mandible may be reduced. *Ptch1, Gli1, Foxd1, Foxd2, Foxf1*, and *Foxf2* are all reported to be the target genes of Hedgehog signaling in cranial NCCs (8). Quantitative RT-PCR analysis showed that the expression of *Shh, Ptch1, Gli1, Foxd1, Foxd2, Foxf1*, and *Foxf2* decreased significantly in the first branchial arch of mutant (Fig. 3C). Wholemount *in situ* hybridization results demonstrated that the expression of Shh was only detected in the proximal epithelium, but much reduced in the distal epithelium of the first branchial arch (Fig 3D, Fig. S4). The expression of *Ptch1* was also down-regulated in the distal region of the mandible of the mutant (Fig. 3D). These results indicated that the deletion of Isl1 in the distal ectoderm caused the down-regulation of the Shh signaling pathway in the distal region of the mandible.

**Figure 3.**
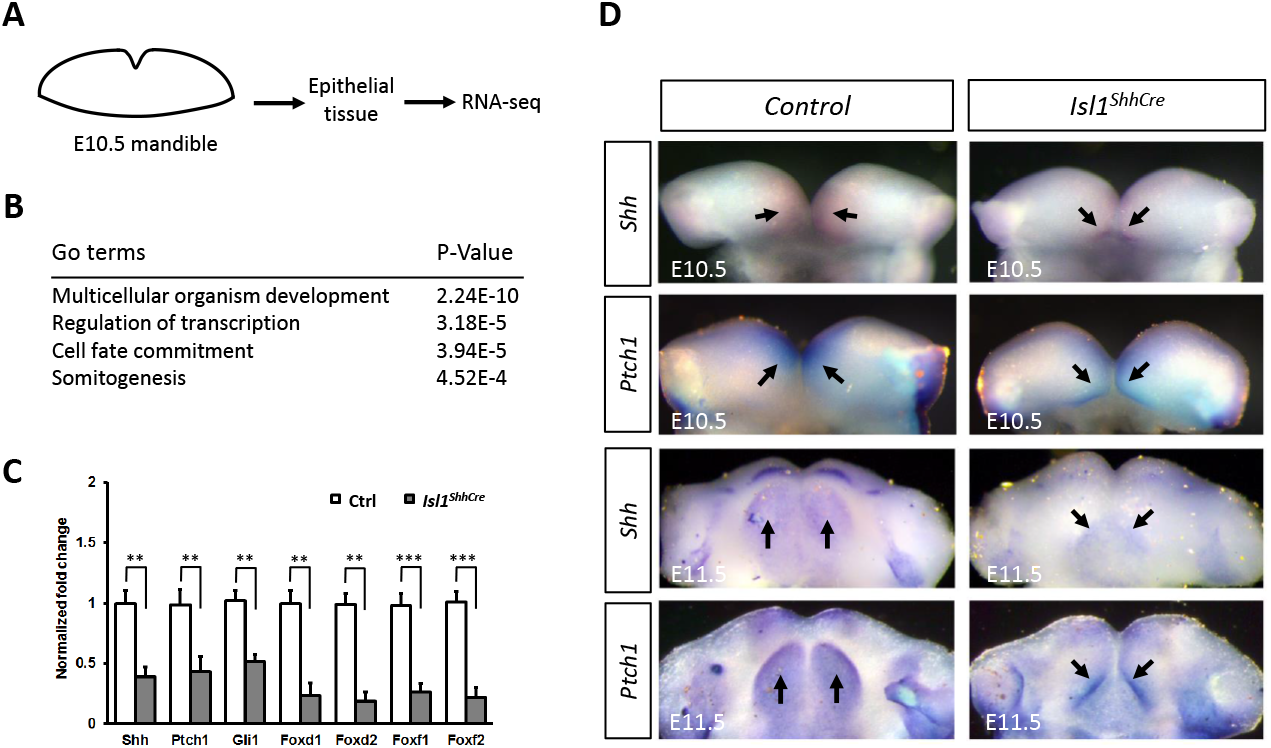
Deletion of Isl1 leads to the down-regulation of Shh in *Isl1*^*ShhCre*^ mandible. (A) Epithelium was isolated from the E10.5 mandible of *Shh*^*Cre*^ (Control) and *Isl1*^*ShhCre*^ mice for RNA-seq analysis. (B) Gene ontology (GO) biological annotation analyses identified high enrichment scores for the categories indicated. (C) Quantitative RT-PCR of *Shh, Ptch1, Gli1, Foxd1, Foxd2, Foxf1*, and *Foxf2* independently in derived mandibular arch RNA samples (n=3 to 6 samples). Error bars represent standard deviations. *, P < 0.05. **, P < 0.01. ***, P < 0.001. Ctrl, Control. (D) Whole-mount *in situ* hybridization showing the expression of *Shh* and *Ptch1* in the *Shh*^*Cre*^ (Control) and the *Isl1*^*ShhCre*^ mutant.

### Downregulation of Shh signaling pathway leads to abnormal expression pattern of CXCL12 in the first branchial arch

It was previously shown that chemokine CXCL12 provides directional cues for migrating CXCR4^+^ muscle progenitor cells in the limb (9). In the developing first branchial arch, CXCL12 is expressed in the mesenchymal cells, and its receptor CXCR4 is expressed in the muscle progenitor cells (9). The co-expression of CXCR4 and Desmin in myoblasts was confirmed by immunofluorescence (Fig. S5). At E10.5, the expression of CXCL12, but not that of CXCR4 was detected in the base of tongue anlage, suggesting that the CXCL12-expressing mesenchymal cells populate the tongue anlage before the invasion of myoblast cells (Fig. 4C, D, C’, D’). At E11.5, the expression domain of CXCL12 and CXCR4 was similarly extended into the middle region of wild type anlage tissue (Fig. 4G, H). In contrast, there is no co-localization of CXCL12 and CXCR4 expression in the *Isl1*^*shhCre*^ (Fig. 4G’, H’), which indicates that CXCR4^+^ myoblasts did not reach the CXCL12 expression domain. In addition, the expression region of CXCL12 in the first branchial arch of the mutant did not extend farther enough, and its expression in the mutant was much higher than that in the wild type (Fig. 4 C, G, C’, G’). CXCL12 expression was only detected in the proximal mesenchyme below the epithelium where Shh is expressed, suggesting that Shh expression in the mandibular epithelium may determine the region where CXCL12^+^ mesenchymal cells located (Fig 4 A, B, C, A’, B’, C’, E, F, G, E’, F’, G’). Thus, we hypothesize that the abnormal expression pattern of Shh and CXCL12 in the first branchial arch led to the migration defect of CXCR4^+^ myoblasts.

**Figure 4.**
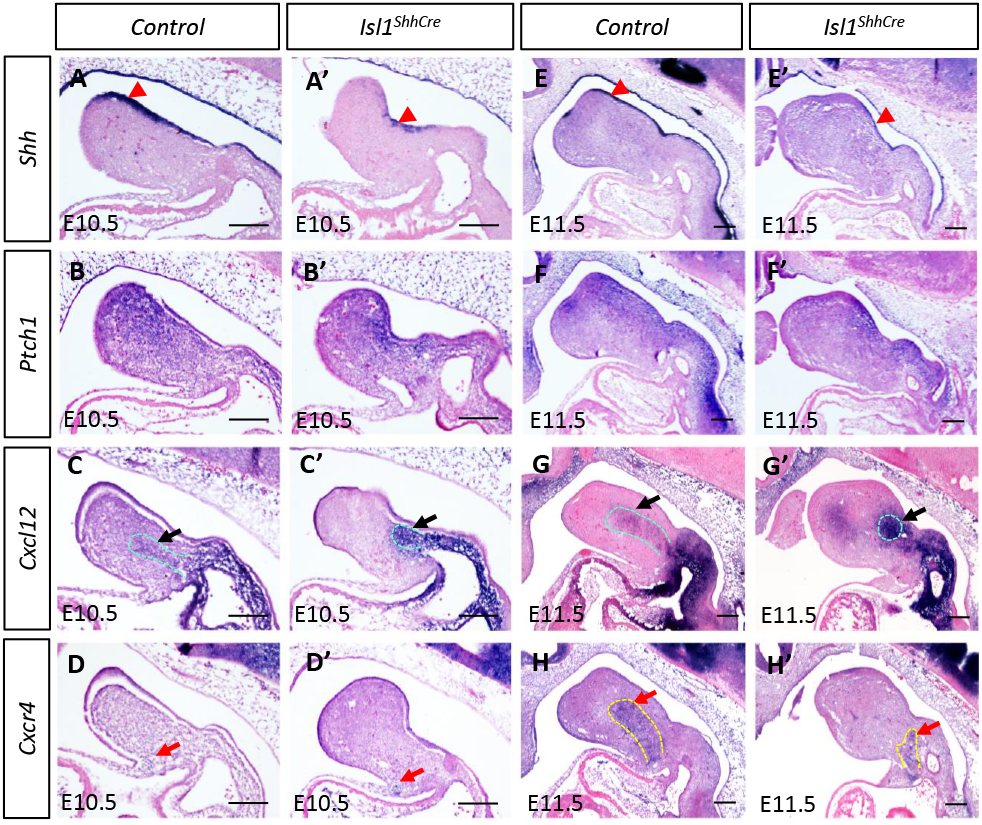
Comparison of Shh and CXCL12 expression in *Isl1*^*ShhCre*^ mandible. *In situ* hybridization of *Shh* (A, A’, E, E’. black arrows), *Ptch1* (B, B’, F, F’. red arrows), *CXCL12* (C, C’, G, G’. red arrowheads), and *CXCR4* (D, D’, H, H’) on the sagittal sections of E10.5 (A, A’, B, B’, C, C’, D, D’) and E11.5 (E, E’, F, F’, G, G’, H, H’) mandibles in *Shh*^*Cre*^ (Control) and *Isl1*^*ShhCre*^ mice. Scale bars, 200 μm.

### Shh regulates the migration of CXCL12^+^ cells through Wnt5a

Our *in situ* hybridization results suggested that CXCL12^+^ cells in the first branchial arch migrated from the adjacent pharyngeal arch. Considering that Wnt5a affects cell migration and can alter outgrowth of many structures including the tongue, it is possible that the reduction of Shh expression limits the migration of CXCL12^+^ mesenchymal cells through Wnt5a. Consistent with this idea, *Wnt5a* expression was detected in the NCCs under the Shh-expressing epithelial cells, but down-regulated in the distal region of the mutant (Fig. 5A). More importantly, Wnt5a expression was strongly induced by Shh-saturated beads (Fig. 5B), or purified recombinant Shh protein in mandible explant culture (Fig 5C). These data suggested that Wnt5a is a downstream target gene of Shh. In addition, Wnt5a-saturated beads could induce the migration of CXCL12^+^ cells (Fig. 5D). The results indicate that the migration of CXCL12^+^ mesenchymal cells is regulated by the Shh/Wnt5a cascade, and the abnormal migration and accumulation of CXCL12^+^ mesenchymal cells in the mutants are due to changes in the expression patterns of Shh and Wnt5a.

**Figure 5.**
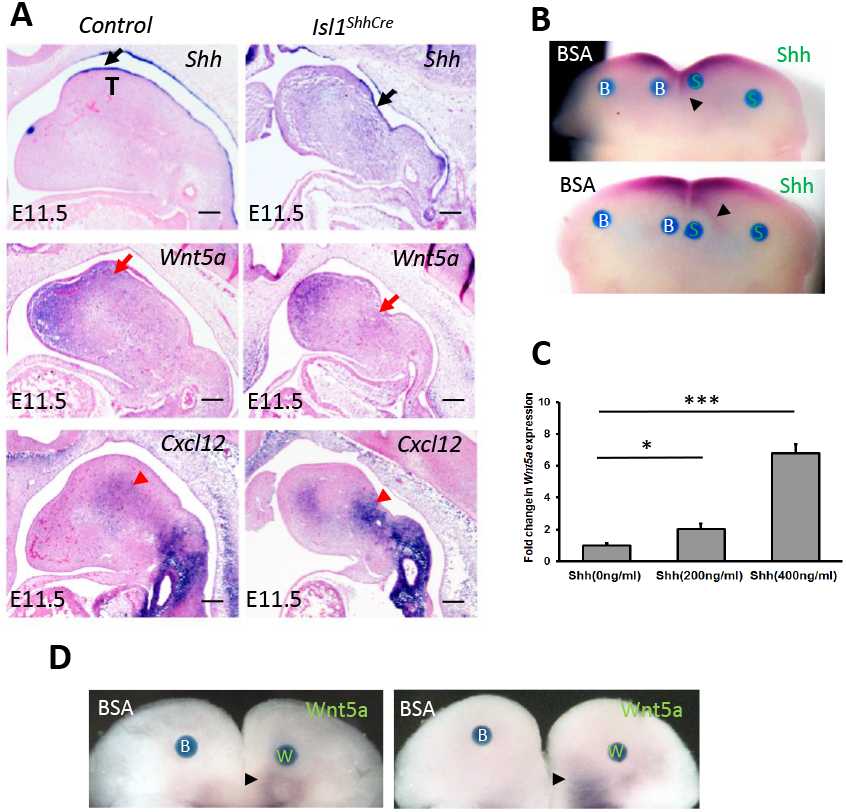
Shh regulates the migration of CXCL12^+^ cells through Wnt5a. (A) In situ hybridization of *Shh* (black arrows), *Wnt5a* (red arrows), and *CXCL12* (red arrowheads) on the sagittal sections of E11.5 mandibles in *Shh*^*Cre*^ (Control) and *Isl1*^*ShhCre*^ mice. (B) Effect of Shh protein on *Wnt5a* expression in mandibular culture. An Shh-saturated, but not bovine serum albumin (BSA)-saturated, bead induces the expression of *Wnt5a* (arrowheads). (C) Quantitative RT-PCR demonstrates the transcription of Wnt5a in mandibular organ cultures. Error bars represent standard deviations (n=3 to 6 samples). Student’s *t* test was used for statistical analysis. *, P < 0.05. **, P < 0.01. ***, P < 0.001. (D) Effect of Wnt5a protein on CXCL12 positive cells migration. A Wnt5a-saturated, but not BSA-saturated, bead attracts the migration of CXCL12 positive cells (arrowheads). B, BSA; S, Shh; W, Wnt5a; T, tongue. Scale bars, 200 μm.

### CXCL12 is a bi-directional cue for myoblast cells

Mandible explant culture was used to determine whether the chemotaxis of myoblasts changes with the concentration of CXCL12. The low concentration (100 ng/μl) of CXCL12 protein saturated beads obviously attracted Desmin^*+*^ myoblast cells to the protein beads (Fig. 6A). On the other hand, Desmin^+^ myoblasts accumulated on a circular band at a certain distance from the high-concentration (500 ng/μl) CXCL12 protein saturated beads (Fig. 6B). The results indicated that high concentration of CXCL12 protein can attract myoblasts to aggregate around, but prevent myoblasts from approaching. The CXCL12 protein diffused from the beads to the surroundings, forming a concentration gradient from high to low. The attraction and repulsion of CXCL12 to myoblasts reached equilibrium when a certain concentration of CXCL12 was reached in the circular band (Fig. 6B).

**Figure 6.**
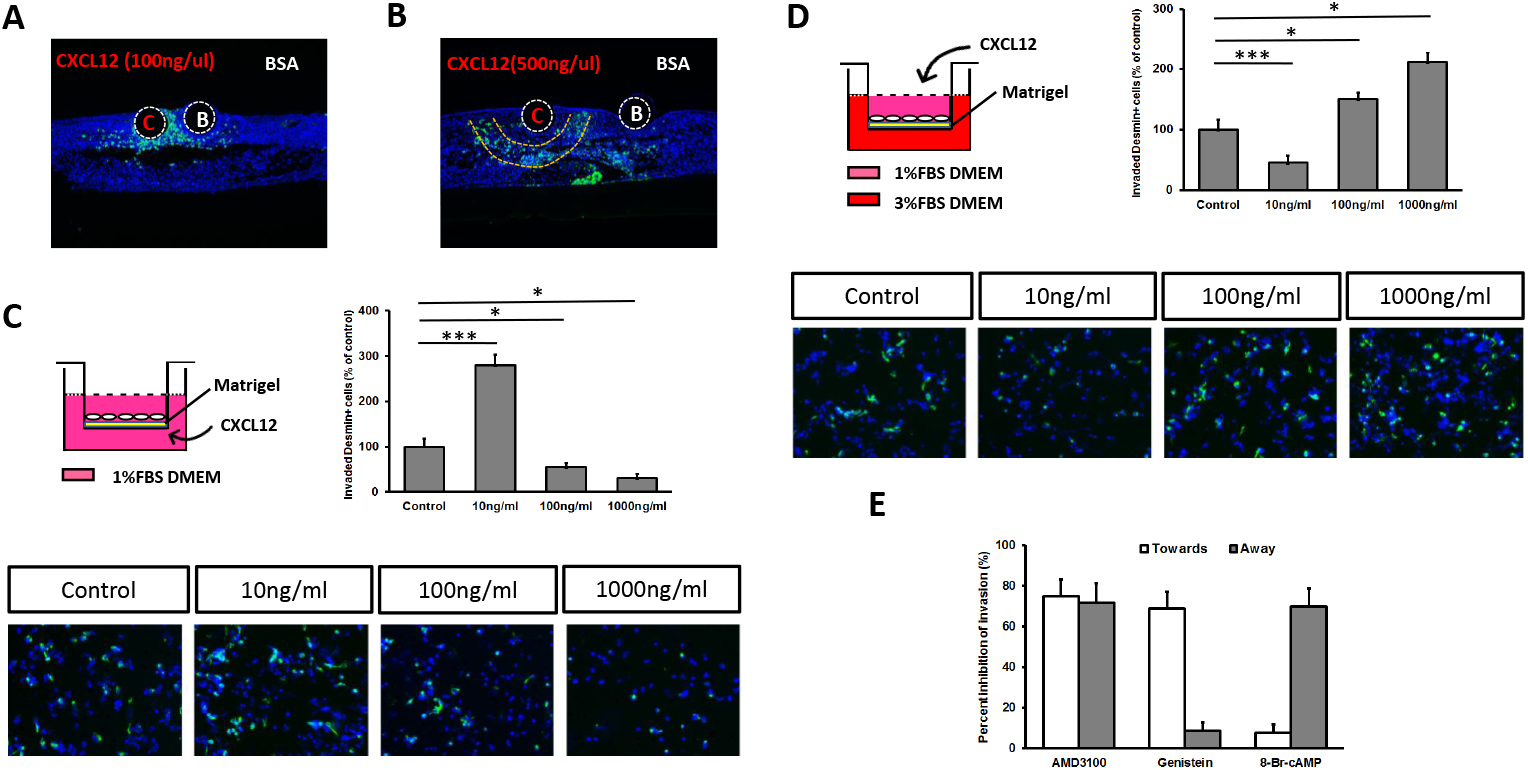
CXCL12 has a concentration-dependent bi-functional chemotactic effect on myoblasts. Immunostaining of Desmin in mandible explants from E11 wild-type mice cultured with BSA control or CXCL12 beads. CXCL12 attracts the migration of Desmin^*+*^ myoblasts at 100 ng/μl (A), but repels them at 500 ng/μl (B). (C) For the chemotaxis assay, primary cells isolated from the E11 tongue tissue of wild-type embryos were added to the upper chamber (2×10 ^5^ per well), and CXCL12 was added to the lower chamber at concentrations (0, 10, 100, and 1000 ng/ml). (D) For the competitive chemotaxis assay, CXCL12 (0, 10, 100, and 1000 ng/ml) was mixed with isolated primary cells (1×10 ^5^ per well) plated to the upper chamber in media containing 1% (v/v) FBS. The media containing 3% (v/v) FBS was added to the lower chamber. The cells on the upper surface of the membrane were removed using a cotton swab after 24 h. The membranes were cut from inserts and mounted onto glass slides for immunofluorescence staining. Desmin antibody was used to label myoblasts. (E) Movement towards and away from CXCL12 has different sensitivities to inhibitors and cyclic nucleotide agonists. Primary cells were incubated with the CXCR4 inhibitor AMD3100, tyrosine kinase inhibitor genistein, and the membrane permeable cAMP agonist 8-BrcAMP, and then were added to the chemotaxis assay. The percent inhibition of movement of Desmin^*+*^ myoblasts towards or away from 10 ng/ml and 1 mg/ml SDF-1 was determined. Error bars represent standard deviations (n=3 to 6 samples). Student’s *t* test was used for statistical analysis. *, P < 0.05. **, P < 0.01. ***, P < 0.001. B, BSA; C, CXCL12.

Primary cultures of a mixture of mesenchymal cells and myoblasts isolated from the E11 tongue tissue of wild-type embryos were used to evaluate the chemotaxis of CXCL12. The inserts were coated with Matrigel matrix to mimic the *in vivo* environment. When CXCL12 was added to the lower chamber, the chemotactic attraction activities of CXCL12 on Desmin^*+*^ myoblast cells were observed with a maximum effect at 10 ng/ml (Fig. 6C). At this concentration, more than 3-fold Desmin^*+*^ cells transmigrated to the lower chamber than that of control. The movement of Desmin^*+*^ myoblasts towards CXCL12 dramatically decreased when CXCL12 concentration was increased to 1000 ng/ml (Fig. 6C). The number of Desmin^*+*^ cells transmigrated was half less than that of the control, which indicated that Desmin^*+*^ myoblasts would move away from CXCL12. The results confirmed that CXCL12 would attract CXCR4^+^ myoblasts at low concentration, but would repulse at high concentration.

Competitive chemotaxis assay was also used to confirm the bidirectional effect of CXCL12 on myoblast invasion. When CXCL12 was added to the upper chamber, the moving away of Desmin^*+*^ myoblasts from CXCL12 reached maximum at 1000 ng/ml, with about 2-fold more Desmin^*+*^ cells transmigrating than that of control (Fig. 6D). Conversely, the number of Desmin^*+*^ cells transmigrated was more than 2-fold less than that of the control when CXCL12 concentration decreased to 10 ng/ml. The data illustrated a concentration-dependent bidirectional movement of myoblast cells in response to CXCL12 (Fig. 6D). The direction of myoblasts invasion was dependent on the concentration of CXCL12, suggesting that CXCL12 functions as an attractant for CXCR4^+^ myoblasts in normal developing tongue tissue, but the abnormally high level of CXCL12 in the primordium of tongue of mutants could prevent myoblasts from invading into it.

Treatment with the selective CXCR4 inhibitor AMD3100 resulted in inhibition of the movement of myoblasts in response to CXCL12 in both directions (away and towards) (Fig. 6E). It is reported that CXCL12 can induce the activation of tyrosine kinase and the inhibition of cAMP (27-29). Pre-incubation of myoblasts with the tyrosine kinase inhibitor genistein only inhibited movement to CXCL12, but had no effect on movement away from CXCL12. In contrast, the membrane permeable cAMP agonist 8-Br-cAMP can block movement away from CXCL12, but has no effect on the movement towards the chemokine. These data indicate that the attraction and repulsion of CXCL12 to myoblasts is mediated by distinct signal transduction pathways.

### Defective tongue development is partially rescued by activation of Hedgehog pathway

To further test the hypothesis that Isl1 regulates tongue development through Hedgehog signaling, we next examined whether Ihh can rescue the defective tongue morphogenesis in *Isl1*^*shhCre*^. In this study, we conditionally expressed another Hedgehog ligand Ihh in the mandibular epithelium using a transgenic allele (Tg-*pmes*-*Ihh*), as Ihh and Shh could activate the same cell signaling pathway (30). As expected, the transgenic *Ihh* indeed alleviated the defect of aglossia, and the *Isl1*^*shhCre*^; Tg-*pmes*-*Ihh* allele displayed microglossia (Fig. 7A, B). The invasion of myoblasts (Pax3^*+*^ and Desmin^*+*^) was rescued after expression of Ihh (Fig. 7C; Fig. S6). Moreover, *Ptch1* expression was induced in the mesenchyme of tongue, indicating the activation of Hedgehog signaling (Fig. 7D). Expression pattern of Wnt5a appeared to become normal, and co-localization of CXCL12^+^ and CXCR4^+^ cells was restored by expression of transgenic *Ihh* (Fig. 6D). Although expression of *Ihh* was insufficient to fully rescue the defect of tongue development, the genetic integration of Hedgehog signaling was evident. The signaling cascade model controlling myoblasts invasion during tongue morphogenesis is diagramed in Figure 8.

**Figure 7.**
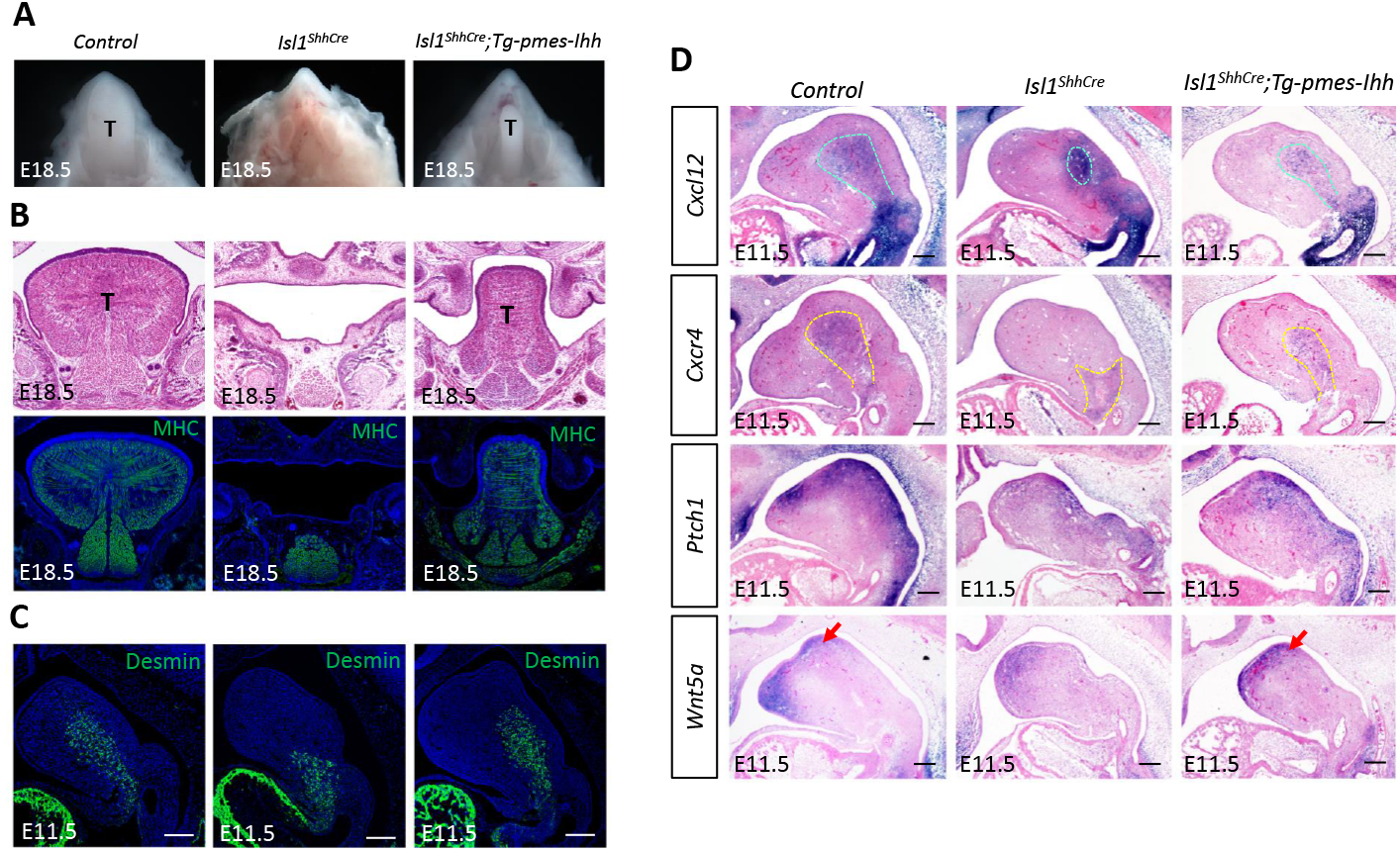
Reactivation of Hedgehog signaling rescues tongue development in Isl1 mutant embryos. (A) Morphological analysis of E18.5 *Shh*^*Cre*^ (Control) and *Isl1*^*ShhCre*^ mice tongue. (B) HE and MHC immunofluorescence staining on the coronal sections of E18.5 mandibles in *Shh*^*Cre*^ (Control) and *Isl1*^*ShhCre*^ mice. (C) Immunofluorescence analysis of Desmin (green) on the sagittal sections of E11.5 *Shh*^*Cre*^ (Control), *Isl1*^*ShhCre*^, *Isl1*^*shh-Cre*^; Tg-*pmes*-*Ihh* mandibles. (D) *In situ* hybridization shows the expression pattern restoration of *CXCL12, CXCR4, Ptch1*, and *Wnt5a* (red arrows) in the *Isl1*^*shhCre*^; Tg-*pmes*-*Ihh* mandibular arch. T, tongue. Scale bars, 200 μm.

**Figure 8.**
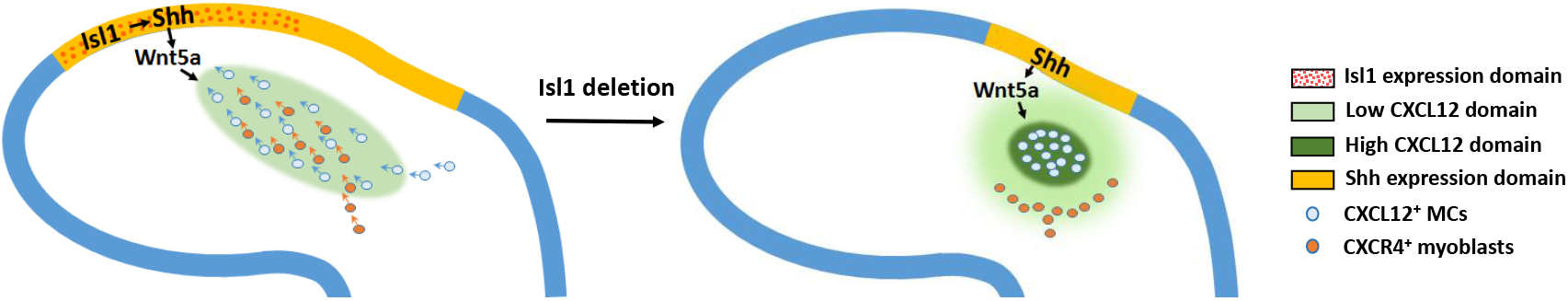
Model of the signaling cascade governing distal myoblast cells invasion in the tongue. In wild-type embryos, Shh is expressed in the mandible distal epithelium cells, and Wnt5a induces the migration of CXCL12 positive cells into the distal mesenchyme domain. CXCR4 positive myoblasts are attracted by CXCL12 and invade into the presumptive tongue primordium. In Isl1 mutant embryos, Shh expression in the mandible distal epithelium cells is missing. CXCL12 positive cells accumulate in the proximal mesenchyme, which leads to high CXCL12 concentration in this domain. The chemotactic repelling effect of high concentrations of CXCL12 leads to the invasion defect of CXCR4 positive myoblasts. MCs, mesenchymal cells.

## Discussion

Although the tongue is an extremely important organ in mammals, the molecular mechanisms of tongue morphogenesis remain poorly understood because of its complex anatomy and cell origins (31). In this study, we demonstrated that transcription factor Isl1 is required for the development of tongue, as the deletion of Isl1 in the epithelium of first branchial arch led to defective invasion of myoblast cells.

Tongue myogenic progenitors originate from the occipital somites and migrate along hypoglossal cord to the tongue primordium at about E10.5 (32). Both Desmin and Pax3 mark the early myogenic precursor cells that migrate from the somites to the branchial arch (25, 26). In this study, the Desmin^+^ and Pax3^+^ myogenic cells were observed in the floor of first branchial arch at E10.5 in both wild type and *Isl1*^*shhCre*^ embryos. However, the myogenic cells did not invade into the tongue primordium after they arrived the first branchial arch in the *Isl1*^*shhCre*^ embryos. It has been reported that NCCs initiate tongue development and populate the tongue primordium prior to the invasion of myogenic progenitors (33). Thus, we hypothesize that the ablation of Isl1 in epithelial cells resulted in a microenvironment in the tongue primordia unconducive to muscle cell invasion. The disconnection between NCC-derived mesenchymal cells and myoblasts in the primordium led to the defective tongue development.

The chemokine receptor CXCR4 and its ligand CXCL12 provide guidance for many different cell types such as lymphocytes, germ cells, hematopoietic cells and muscle progenitors (16-20). All these different cell types are accurately guided by CXCL12 to different targets at the same time, which suggests tight control of the spatial and temporal distribution of CXCL12 (34). CXCL12 is expressed in the mesenchyme of limb bud and the first branchial arch, whereas its receptor CXCR4 is expressed in migrating muscle progenitors (9). However, CXCR4 positive muscle progenitor cells did not migrate to the center of the limb bud where CXCL12 was highly expressed, but were distributed around (9). It highly suggests that the expression level of CXCL12 might regulate the distribution of muscle progenitor cells. Here, we found abnormal high CXCL12 mRNA expression in the mandibular arch of *Isl1*^*shhCre*^ embryos. We propose that this higher CXCL12 expression in the presumptive tongue primordium lead to the invasion defect of myoblasts. In keeping with this notion, our chemotaxis assay demonstrated that CXCL12 had a concentration-dependent bi-functional effect on myoblasts invasion, attracting CXCR4^+^ myoblasts invasion at low concentration but repelling at high concentration. The invasion and distribution of CXCR4^+^ myoblasts during tongue and limb development may be precisely regulated by the intensity of CXCL12 expression. This is the first study to discover that CXCL12 has a bi-functional effect on myoblast invasion. In agreement with our observations, a previous study reported the similar bi-functional effect of CXCL12 in T cell migration (35).

How might Isl1 in the distal ectoderm alter the expression of CXCL12 in the mesenchyme? Down-regulation of Shh in the distal epithelium after Isl1 ablation is associated with altered CXCL12 expression and the impaired migration of CXCL12^+^ mesenchymal cells. Our data indicates that Wnt5a is a target of Shh, which is consistent with the report in hair follicle morphogenesis (36). Several lines of evidence suggest that Wnt5a may mediate Shh signaling in the regulation of CXCL12^+^ mesenchymal cells migration. First, it is well documented that Wnt5a plays a critical role in regulating cell migration and polarity (10). Second, the expression of Wnt5a is lost in the tongue of *Isl1*^*shhCre*^ embryos. Conversely, activation of SHH signaling in tissue explants can induce Wnt5a expression. Third, Wnt5a attracts the migration of CXCL12^+^ mesenchymal cells in the first branchial arch. Moreover, the partial restoration of tongue morphogenesis and the adequate recovery of myoblast invasion by expression of transgenic Ihh further provide strong support that Hedgehog signaling is critical for tongue development. Based on these findings, we propose that Isl1 regulates the migration of CXCL12^+^ mesenchymal cells through the Shh/Wnt5a signaling cascade. In *Isl1*^*shhCre*^ mutants, CXCL12^+^ cells clustered in a relatively small proximal region of the mutant mandibular arch, leading to a high CXCL12 concentration in this domain. The distal migration of CXCL12 expressing cells is essential for establishing the concentration gradient of CXCL12, which attracts myoblast invasion during tongue development.

The binding of CXCL12 to its receptor CXCR4 triggers multiple intracellular signaling pathways related to chemotaxis (27-29). It has been reported that both tyrosine kinase and cyclic nucleotides are essential for cell polarization and migration (37-40). Our data suggests that the attraction of CXCL12 to myoblasts is mediated by the activity of tyrosine kinase, while cAMP is responsible for the repulsion of CXCL12 to myoblasts. It is possible that different concentrations of CXCL12 activate the distinct signal transduction pathways downstream of its receptor CXCR4, resulting a ligand concentration-dependent bi-functional effect on myoblast invasion. In addition to the regulation of migration processes, CXCL12/CXCR4 signal pathway also promotes growth and survival of different cells types (41-43). Consistently, defective proliferation of myoblasts was detected at E11.5. The reduced cell proliferation might be due to the lack of interactions between CXCL12^+^ cells and CXCR4^+^ myoblasts in the mutant embryos. However, the process of differentiation of the myoblasts was not affected, indicating that the reciprocal interactions between these two types of cells are necessary for the myoblasts proliferation but not differentiation, in keeping with the previous finding that myoblast differentiation was not affected after CXCR4 inhibitor treatment (21).

To the best of our knowledge, this is the first study to report that Isl1/Shh/Wnt5a axis regulates CXCR4^+^ myoblast invasion by regulating the migration of CXCL12 expressing mesenchymal cells. The concentration-dependent bidirectional regulation of CXCL12 on the migration and invasion of CXCR4^+^ myoblast may also be significant for the development of other organs such as limbs. Our results will promote the understanding of the mechanism of myoblast invasion and open perspectives for the pathogenesis study of human tongue deformities and related syndromes.

## Materials and Methods

### Animals

All animal procedures were approved by Hangzhou Normal University Animal Care and Use Committee (HNLA-2012-018). Construction of conditionally targeted *Isl1*^*fl*^ mice, *Isl1*^*LacZ*^, and Tg-*pmes-Ihh* mice has been described previously (24, 44). *Shh*^*tm1(EGFP/cre)*^ and *R26R-LacZ* were purchased from the Jackson Laboratory and maintained on a C57BL/6J background. The morning of observed vaginal plug was designated as day 0 (E0) of pregnancy.

### Scanning electron microscopy analysis

Embryos were fixed in 2.5% glutaraldehyde in 0.1 M phosphate buffer (pH 7.4) overnight. After rinsing in PBS the sections were post fixed with 1% osmium tetroxide solution for 1.5 h. Dehydrate samples using a series of acetone solutions: 30%, 50%, 70%, 80%, 90%, 100% (three times) for 15 min each. Samples were critically point-dried after a 15 min treatment with isoamyl acetate. Then the samples were mounted on conductive paper and sputter-coated with gold. Images were recorded with a scanning electron microscope (Hitachi S-3000N) with a 15-kV accelerating voltage.

### X-gal staining

X-gal staining of whole-mount samples was performed as described previously (24). Embryos were fixed in 4% PFA solution (5mM EGTA and 2mM MgCL2) at 4°C for 1 h. The fixed embryos were rinsed three times in washing buffer (2mM MgCL2, 0.01% sodium deoxycholate, and 0.02% NP40 in PBS). The embryos were then incubated in staining solution (0.1% X-gal, 5 mM potassium ferricyanide, 5 mM potassium ferrocyanide, 2 mM Tris (pH 7.3) in washing buffer) overnight in the dark at 37°C. Lastly, the stained embryos were washed in PBS and used for whole mount analysis. The X-gal stained sections were counterstained with Nuclear Fast Red.

### RNA-seq analysis

Mouse mandibular epithelial tissue were carefully collected from E10.5 wild-type or *Isl1*^*shhCre*^ embryos. Total RNA was isolated from epithelial tissues as previously described (24, 45). RNA-seq libraries were established with an Illumina TruSeq RNA sample prep kit and sequenced using Illumina HiSeq 4000 with 150 bp paired-end sequencing strategy. Expression analysis was performed using HISAT2 and DESeq2 (46, 47), and differentially expressed genes were determined with the cutoff fold change≥1.5 and P-value ≤ 0.05. Function enrichment analysis was performed with the DAVID tools (https://david.ncifcrf.gov/). All the raw data generated from RNA-seq in this work was deposited in BIGD (bigd.big.ac.cn) under the accession number CRA004944.

### Quantitative realtime-PCR

Total RNA from mouse mandibles was extracted from organ culture explants using TRIzol Reagent (Thermo Fisher Scientific). All RNA samples were treated with DNA-free DNA Removal Kit (Thermo Fisher Scientific). The first-strand complementary DNA (cDNA) was then synthesized from 1μg of total RNA from each sample using a PrimeScript 1st Strand cDNA synthesis kit (TaKaRa). Real-time RT-PCR was performed using the following primer sequences: *Shh*, 5’-AAAGCTGACCCCTTTAGCCTA and 5’-TGAGTTCCTTAAATCGTTCGGAG; *Ptch1*, 5’-AGACTACCCGAATATCCAGCACC and 5’-CCAGTCACTGTCAAATGCATCC; *Gli1*, 5’-CCAAGCCAACTTTATCAGGG and 5’-AGCCCGCTTCTTTGTTAATTTGA; *Foxd1*, 5’-AAAATCGCCCTATGCTGC and 5’-CTGGACCTGAGAATCTCTACACC; *Foxd2*, 5’-ATTTATGAAGAGTCTCCAGACC and 5’-GATGCTCAAACAGAAAAGC; *Foxf1*, 5’-ACTCCAGTGTCTTTCACCTTGC and 5’-TGAGCCTGAACTACACCAGC; *Foxf2*, 5’-TCAGTAGGACATTTCTTCC and 5’-CTGTCACAATACTGAGAGC; *Wnt5a*, 5’-CTAACAAGTGTGACAAGATCC and 5’-CTTGAAAGCAATGTCTAGC; *18S*, 5’-TAGAGGGACAAGTGGCGTTC and 5’-CGCTGAGCCAGTCAGTGT. The specificity for each primer set was confirmed by electrophoresis and sequencing before using. The real-time PCRs were performed in triplicate using SsoFast EvaGreen Supermix as previously described (24). The relative amount of each gene transcript was calculated using the 2^-ΔΔCT^ method and normalized to the endogenous reference gene *18S*. Data were analyzed with CFX Manager software and are represented as means ± standard errors of the means (SEM).

### Immunohistochemistry and immunofluorescence

The collected embryos were fixed in 4% paraformaldehyde solution, dehydrated, embedded in paraffin and sectioned as previously described (48). Heat induced epitope retrieval (HIER) was used to unmask antigen epitopes. For the cultured cells, 4% paraformaldehyde (PFA) solution was used for 10 min. After blocking in 1% bovine serum albumin, the sections were incubated with antibodies against MHC (Sigma), Desmin (Abcam), Cylin D1 (Abcam), Pax3 (Abcam), or Ki67 (Abcam) at 4°C overnight. Then, HRP conjugated secondary antibody against the primary antibody was applied for 1h at room temperature. For immunofluorescence, secondary antibodies conjugated with Alexa Fluor 488 or 594 (1:1000, Invitrogen) were used. Images were analyzed using a microscope Leica DM4 B equipped with a digital camera.

### Histological analysis and in situ hybridization

Whole-mount and section *in situ* hybridization was performed as previously described (24). For section in situ hybridization, samples were fixed in 4% paraformaldehyde solution, dehydrated, embedded in and sectioned at 12 μm. For whole-mount in situ hybridization, samples were fixed in freshly made 4% paraformaldehyde/PBS, dehydrated into methanol and bleached with 6% hydrogen peroxide (H2O2). Non-radioactive antisense RNA probes were generated by *in vitro* transcription using DIG RNA labeling kit (Roche Diagnostics). For histological analysis sections were stained with Hematoxylin and Eosin according standard protocols.

### Cell proliferation assay

Cell proliferation activity was evaluated by 5-bromodeoxyuridine (BrdU) labeling and immunofluorescence staining. Timed pregnant mice were sacrificed 30 min after after a single intraperitoneal injection of BrdU (Sigma) at 3 mg/100 g body weight. The collected embryos were fixed in 10% neutral buffered formalin at 4°C overnight and embedded in paraffin. BrdU-labeled cells were detected immunohistochemically in paraffin sections according to the manufacturer’s instructions. The cell growth ratio (%) was calculated by dividing the number of BrdU positive nuclei by the total number of nuclei.

### Chemotaxis assay

The chemotactic response of mandible myoblasts to CXCL12 was evaluated using a 24-well fitted transwell inserts with membranes (8 μm pore size, Corning Falcon). The inserts were then coated with Corning Matrigel matrix (Fisher scientific) at 200 μg/ml. Mouse tongues were carefully dissected from E11 wild-type embryos. To remove the epithelium, the explants were incubated in 2.25% trypsin/0.75% pancreatin on ice for 5 min. One hundred microliters of isolated primary cells (mixture of myoblasts and mesenchymal cells) were added to the upper well of the chamber (2×10^5^ per well). Both upper and lower chambers were loaded with DMEM containing 1% fetal bovine serum (FBS). CXCL12 (R&D Systems, Minneapolis, MN) was used at concentrations of 0 ng/ml, 10 ng/ml, 100 ng/ml, and 1000 ng/ml in the lower chambers of the transwell. For the inhibitor treatment, cells were pretreated with 0.5 μg/ml AMD3100 (30 min at 37 °C), 1 μg/ml genistein (20 min at 37 °C), or 100 μM 8-Br-cAMP (15 min at room temperature) before added to the upper well of the chamber in selected experiments. For the competitive chemotaxis assay, CXCL12 (0, 10, 100, and 1000 ng/ml) was mixed with isolated primary cells (1×10^5^ per well) plated to the upper chamber in media containing 1% (v/v) FBS. The media containing 3% (v/v) FBS was added to the lower chamber. The cells on the upper surface of the membrane were removed using a cotton swab after treatment for 24 h. The membranes were cut from inserts and mounted onto glass slides for immunofluorescence staining. Desmin antibody was used to label myoblasts. All experiments were repeated for 3-6 times, and the number of invaded Desmin^+^ myoblasts was calculated.

### In vitro organ culture and bead implantation

For bead implantation experiments, mouse mandibles were carefully dissected from E10.5 wild-type embryos and placed into Dulbecco’s modified Eagle medium (DMEM; Gibco) supplemented with 10% FCS. The explants were placed on a Nuclepore Track-Etch membrane (0.2-μm pore size) in Trowel-type organ culture dishes. BSA (1μg/μl), Wnt5a-saturated agarose beads (0.5 μg/μl), CXCL12-saturated agarose beads (0.1 and 1 μg/μl) or Shh-saturated agarose beads (1 μg/μl) were grafted in the mandible. After being cultured for 16 h in a humidified atmosphere of 5% CO2 at 37°C, the explants were fixed in 4% PFA overnight at 4 °C, washed in PBS and finally stored in methanol at - 20 °C until analysis by whole-mount in situ hybridization. Wnt5a, CXCL12 and Shh were purchased from R&D Systems (Minneapolis, MN).

### Statistical analyses

All data are presented as means ± SEM. Student’s t test was used to compare the differentials between data sets. The threshold for statistical significance was P < 0.05.

## Acknowledgments

This work was supported by grants from the National Natural Science Foundation of China (81871166, 81670971 to F. L.) and Natural Science Foundation of Zhejiang Province (LY21C120002 to J. L., Y20C120004 to D. Y.).

## Supplementary figure legends

**Figure S1.**
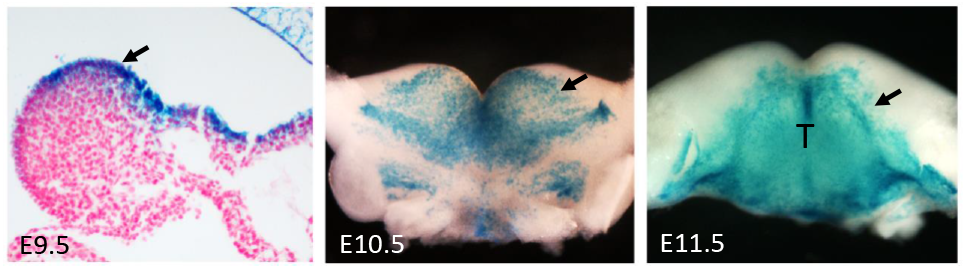
X-Gal staining showing Shh-Cre activity in E9.5, E10.5 and E11.5 *Shh*^*Cre*^ /*R26R* mandibular arch (arrows). T, tongue.

**Figure S2.**
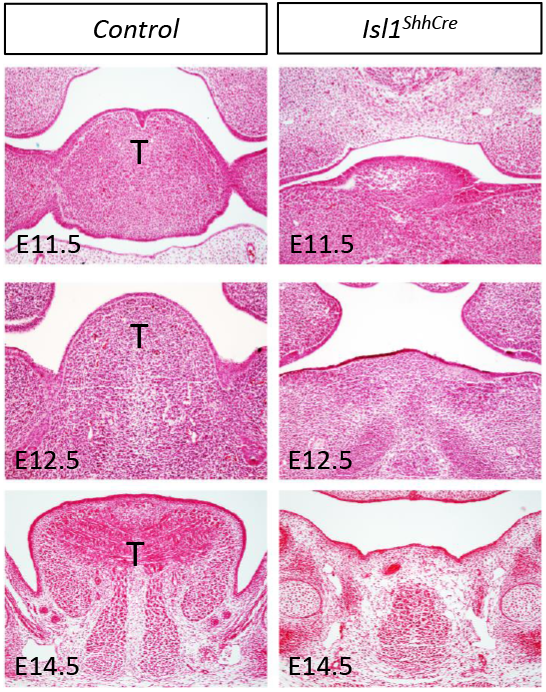
Hematoxylin and eosin staining on the coronal sections of mouse heads showing the tongue of *Shh*^*Cre*^ (Control) and *Isl1*^*ShhCre*^ mice. Deletion of Isl1 in the epithelial cells leads to the defect of tongue development. T, tongue.

**Figure S3.**
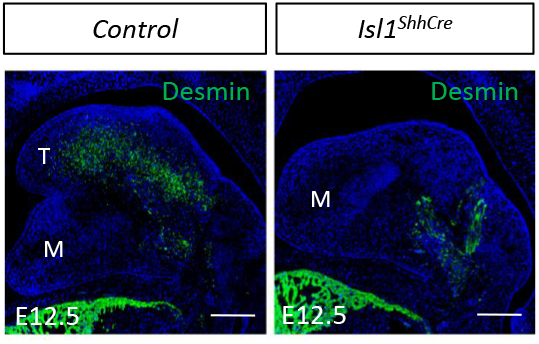
Deletion of Isl1 in the epithelial cells leads to the defect of myoblast cells invasion. Immunofluorescence analysis of Desmin (green) distribution in the tongue of E12.5 *Shh*^*Cre*^ (Control) and *Isl1*^*ShhCre*^ embryos. M, mandible; T, tongue; Scale bars, 400 μm.

**Figure S4.**
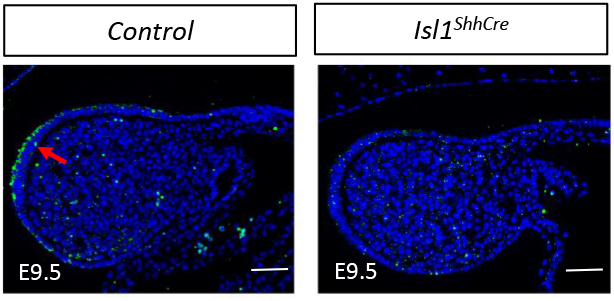
The expression of Shh protein significantly decreased in the *Isl1*^*ShhCre*^ mandibles. Immunofluorescence analysis of Shh protein (red arrows) on the sagittal sections of E9.5 *Shh*^*Cre*^ (Control) and *Isl1*^*ShhCre*^ embryos. Scale bars, 200 μm.

**Figure S5.**
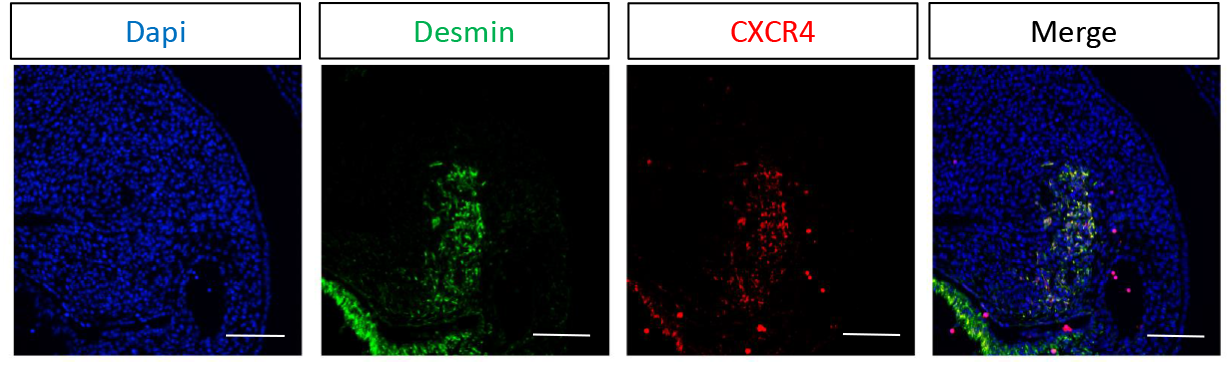
CXCR4 and Desmin are coexpressed in the myoblasts in the mandible. Double immunofluorescence staining with Desmin (green) and CXCR4 (red) on the sagittal sections of E11.5 mandibles. Scale bars, 200 μm.

**Figure S6.**
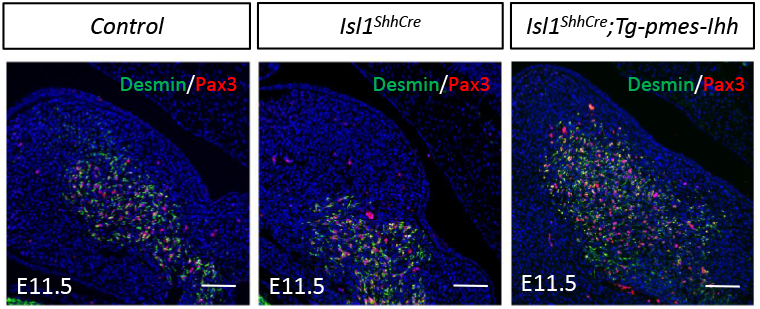
Reactivation of Hedgehog signaling rescues myoblast cells invasion in *Isl1*^*shh-Cre*^ embryos. Double immunofluorescence staining with Desmin and Pax3 antibody on the sagittal sections of E11.5 *Shh*^*Cre*^ (Control), *Isl1*^*ShhCre*^, *Isl1*^*shh-Cre*^; Tg-*pmes*-*Ihh* mandibles. Scale bars, 200 μm.

